# Nucleophosmin mutations lead to abnormal, but reversible, nucleoli architecture and aggregate formation – Implications for NPM1-targeting therapies in AML

**DOI:** 10.1101/2024.12.30.630786

**Authors:** Martin Grundy, Kellie Lucken, Xiaomeng Xing, Eva L. Simpson, Ahmed Bayyoomi, Alison J. Beckett, Ian A. Prior, Daniel G. Booth, Claire H. Seedhouse

**Author notes:** **Corresponding authors**: Dr Martin Grundy, Dr Claire Seedhouse, Dr Daniel Booth.

## Abstract

Mutations in the *NPM1* gene represent the most common (>30% of patients) genetic alteration in Acute Myeloid Leukaemia (AML) and results in the mis-localisation of the mutated NPM1 protein from a predominantly nucleolar localisation to a predominantly cytoplasmic distribution. Numerous studies of NPM1 mutated AML have focussed on the aberrant cytoplasmic localisation of the mutated protein but efforts to reverse this mis-localisation therapeutically have so far resulted in limited clinical benefit. More recently, attention has shifted towards the nucleus with studies showing that mutant NPM1 binds to specific chromatin regions, where it directly regulates oncogenic gene expression. Here, we use high resolution imaging to demonstrate that Nucleophosmin (NPM1) is critical for maintaining normal nucleoli architecture and specifically the integrity of the nucleoli rim. We report for the first time that NPM1 mutated cell lines and primary samples have aberrant nucleoli architecture and demonstrate that the abnormal nucleoli phenotype is reversible. We also report the novel finding that NPM1 mutated protein forms distinct aggregates in NPM1 mutated cells and characterise these for the first time. This work reveals how nucleolar organisation contributes to the molecular mechanisms underpinning NPM1 driven AML and reveals unexpected novel vulnerabilities to be exploited for therapeutic intervention.

## Introduction

Acute myeloid leukaemia (AML) is characterised by the clonal proliferation of undifferentiated myeloid blast cells which can infiltrate the bone marrow, blood and other tissues leading to ineffective haematopoiesis, bone marrow failure and severe cytopenia’s. Although people of all ages can develop AML it is generally a disease of the elderly with a median age at diagnosis of 68 years [1]. Despite advances in supportive care and relapse monitoring, prognosis remains dismal, with only 9.4% of patients who are 65 years and older at diagnosis achieving a 5-year survival [1, 2]. Unlike many cancers, the prognosis for AML has shown little improvement in over 20 years. There is clearly an unmet need to better understand the molecular mechanisms that trigger and drive leukaemia in these patients, providing a tractable approach for discovery of new therapeutic targets. Mutations in the Nucleophosmin (*NPM1*) gene represent the most common genetic alteration in AML occurring in around 35% of adult AML [3]. NPM1 is a ubiquitously expressed and functionally diverse phosphoprotein with reported roles in ribosome biogenesis and transport, DNA repair, centrosome duplication and as a protein chaperone [4–6]. Correct subcellular localisation of NPM1 is critical for normal cellular homeostasis. NPM1 routinely shuttles between the cytoplasm and nucleus [7], but under normal physiological conditions NPM1 localises predominantly to the nucleolus [8], the largest sub-compartment of the nucleus.

First observed more than 200 years ago, nucleoli are membrane-less subnuclear compartments that are the cellular site for ribosome synthesis and assembly and additionally comprise proteins involved in stress response and cell cycle regulation [9]. The lack of enclosing membrane allows nucleoli to respond dynamically to cellular signals by changing their size and protein composition. Nucleoli form around pre-ribosome RNA gene repeats, termed nucleolar organiser regions (NORs). In humans, NORs are located on the short arms of acrocentric chromosomes 13, 14, 15, 21 and 22 [10]. The nucleolus, which is formed by liquid-liquid phase separation, has long been considered as having three droplet-like layers of varying miscibility: the fibrillar centre (FC), the dense fibrillar component (DFC) and the granular component (GC). More recently, the comprehensive spatiotemporal mapping of nucleolar architecture, revealed a fourth nucleoli sub-compartment, the nucleoli rim, and reported that 157 nucleoli proteins, including NPM1, had a characteristic nucleoli rim-like localisation [11].

*NPM1* insertion mutations are AML specific and almost always occur at exon 12, causing a frameshift in the C-terminus encoding region which hampers the correct folding of NPM1 [12]. The altered reading frame leads to loss of the nucleolar localisation signal (NLS) and generation of an additional nuclear export signal (NES), thought to increase binding to the nuclear export protein XPO1, ultimately resulting in the aberrant cytoplasmic distribution of the mutated protein [13–15]. More than 50 different mutations in the *NPM1* gene have been identified all of which result in mis-localisation of the mutated protein (denoted NPM1c+) to the cytoplasm, [16]. This cytoplasmic mis-localisation of NPM1c+ also leads to *HOX* gene upregulation which is critical for leukemogenesis and is required for disease maintenance [17]. Most studies of NPM1 mutated AML have focussed on the aberrant localisation of the mutated protein from the nucleolus to the cytoplasm, along with factors that are expelled from the nucleus to the cytoplasm in consort, including CTCF, FBW7γ and PU.1 [18–20]. This includes efforts to reverse NPM1c+ mis-localisation with SINE (selective inhibitor of nuclear export) compounds such as Selinexor and Eltanexor, but these have so far resulted in limited clinical benefit [21, 22]. More recently, attention has shifted towards the nucleus with studies showing that mutant NPM1 binds to specific chromatin regions, where it directly regulates oncogenic gene expression [23, 24]. Due to stalled progress in development of therapeutic targets for AML and reflecting on new advances mapping nucleolar architecture, we identified a clear need to revisit the molecular dynamics of NPM1 in the context of AML. Using high resolution imaging we performed a comprehensive survey of wild-type (wt) and mutant NPM1 subcellular distribution, revealing multiple novel features, including the role of NPM1wt in maintaining normal nucleoli architecture and demonstrate loss of nucleoli rim structure and distorted nucleoli following NPM1wt knockdown. We report for the first time that this aberrant nucleoli phenotype is also present in NPM1 mutated cell lines and primary samples and demonstrate that the abnormal nucleoli phenotype is reversible. Finally, we also report the intriguing finding that NPM1 mutated protein forms distinct aggregates in NPM1 mutated cells and perform the first characterisation of these enigmatic structures. This work not only contributes to our understanding of the molecular mechanisms underpinning NPM1 driven AML but also reveals an unexpected novel vulnerability to be exploited for therapeutic intervention.

## Results

### NPM1 wild-type knockdown results in loss of the nucleoli rim and nucleoli distortion

Previously, some studies have reported that NPM1 depletion can cause perinucleolar heterochromatin rearrangement (and the disruption of epigenetic marks) [25], and others that NPM1 depletion results in loss of general nucleolar structure [25, 26]. However, confirmation of NPM1 distribution, function and consequences of manipulation (via depletion or expression of mutants), explicitly relating to the nucleolar rim, remains elusive. To address this, we conducted targeted NPM1 depletion using siRNA followed by microscopic analysis. Western blotting and band densitometry (not shown) revealed a 99% reduction in NPM1 abundance following a 6-day RNAi period (Figure 1A). Importantly, expression levels of nucleolin, another nucleoli rim localising protein [11], remained unchanged, underscoring the specificity of NPM1 knockdown (Figure 1A). Immunofluorescence imaging confirmed clear NPM1 and nucleolin localisation at the nucleoli rim (Figure 1Bi yellow arrows). Neither protein was present at the nucleolar rim following 6 days of NPM1 RNAi. Instead, knockdown of NPM1 resulted in distorted nucleoli and a diffuse staining pattern of both nucleolin and the residual NPM1 within the remaining nucleolar space (Figure 1Bii red arrows). This suggests that NPM1 has a pivotal role in maintaining nucleoli rim structure. Next, we performed rescue experiments to test whether loss of nucleolar integrity was reversible. Following 2 days of NPM1 RNAi, cells were transfected to express a scarlet-tagged version of NPM1wt (Scar_NPM1_wt). Remarkably, re-expression of NPM1wt resulted in almost complete correction of normal nucleolar phenotype (Compare figure 1Civ blue arrows with Figure 1Ci yellow arrows), confirming the essential role of NPM1 in maintaining normal nucleoli rim structure. Interestingly, experimental controls revealed that ectopic over expression of NPM1 (on a control RNAi background), regularly caused the formation of large single nucleoli, but that even in this instance NPM1wt was still mostly detectable at the nucleolar rim (Figure 1Cii yellow arrow). This work supports previous studies that there is likely a stringent cellular tolerance of NPM1 and that normal NPM1 function is tightly coupled to appropriate abundance.

**Figure 1.**
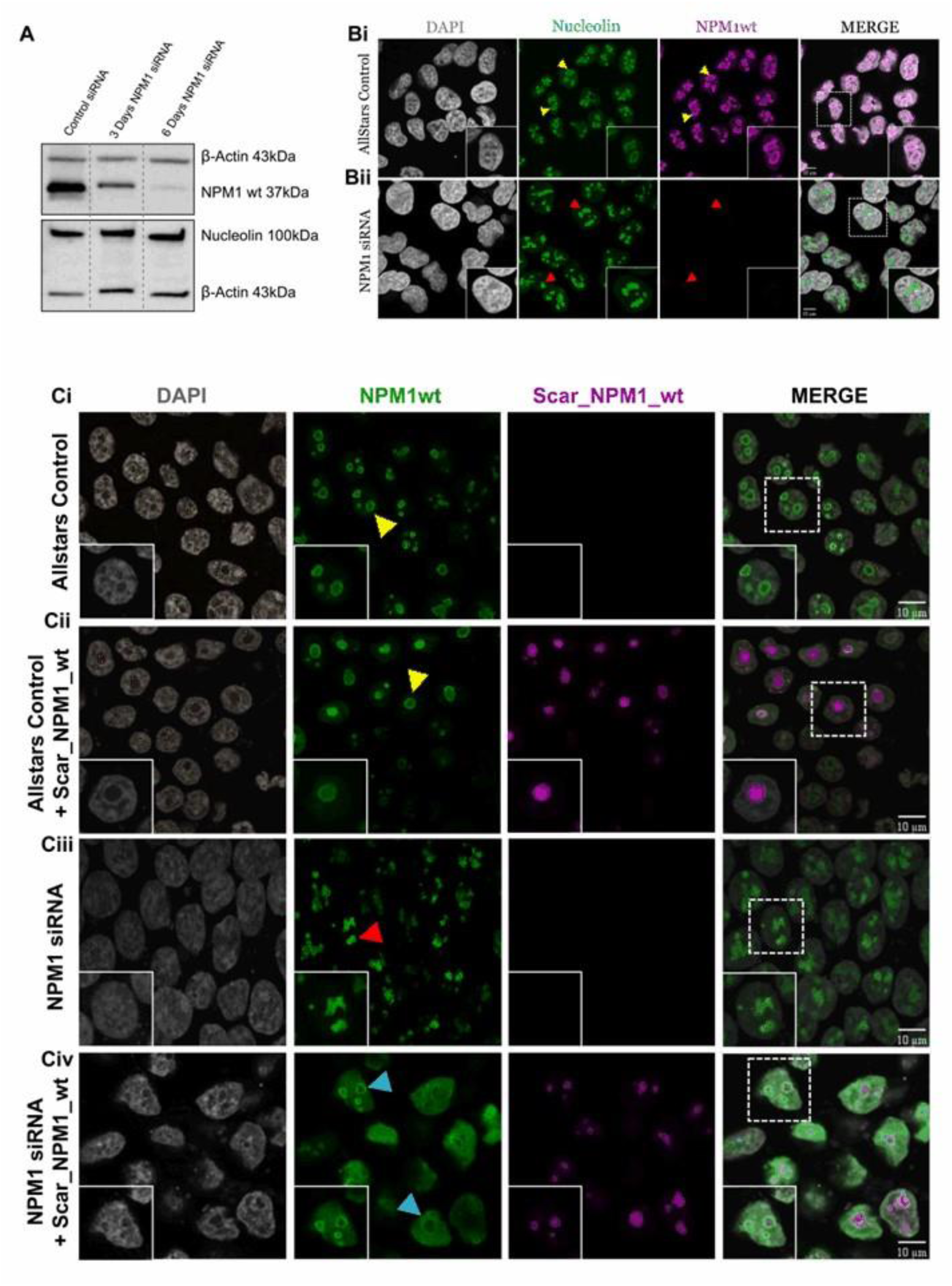
NPM1 wild-type knockdown results in loss of the nucleoli rim and nucleoli distortion. **(A)** NPM1wt and nucleolin protein expression in Hela cells following 3 or 6 days NPM1wt siRNA interference. **(B)** Hela cells were stained with DAPI and antibodies against NPM1wt and nucleolin following 6 days NPM1wt siRNA interference or AllStars negative control. Normal NPM1wt and nucleolin nucleoli rim staining (yellow arrows). Cells lacking a nucleoli rim with distorted nucleoli (red arrows). **(C)** HEK-293T cells were subjected to 2 days NPM1wt siRNA interference followed by 24 hrs transfection with Scar_NPM1_wt and then stained with DAPI and NPM1wt antibody. Normal NPM1wt nucleoli rim localisation (yellow arrows) and distorted nucleoli following NPM1wt siRNA interference (red arrows). Cells with normal NPM1wt nucleoli rim localisation following Scar_NPM1_wt ectopic rescue (blue arrows). All imaging was performed using a Leica DMI4000 B confocal microscope.

### Subcellular localisation of wild-type and mutant NPM1 revealed by high-resolution confocal microscopy

Next, to determine the impact of NPM1 mutation on nucleolar architecture, we closely examined the subcellular localisation patterns of both NPM1wt and the NPM1c+ mutant, using high resolution confocal microscopy. Initially, we chose a simplified system expressing scarlet tagged NPM1wt (Scar_NPM1_wt) or emerald tagged NPM1c+ mutant (Em_NPM1_mut) in both HEK-293T cells (Figure 2A) and HeLa cells (Figure 2B). As expected, NPM1wt localised to the nucleoli (yellow arrows) whilst NPM1c+ predominantly localised to the cytoplasm (red arrows) in both cell types (Figures 2A and 2B). Interestingly, a subtle difference was observed in the nucleolar-to-cytoplasmic distribution of NPM1C+, between the two cell types. In HEK-293T cells NPM1c+ was almost entirely cytoplasmic, whereas in HeLa cells a significant fraction (17.8%) localised to the nucleolus (Figure 2C). This cell type variability is consistent with previous reports, suggesting that the discrepancy might be due to higher endogenous NPM1 expression in HeLa cells compared with HEK-293T cells [27]. Co-transfection (Scar_NPM_wt + Em_NPM1_mut) resulted in increased presence of NPM1 mutant protein in the nucleoli of both HEK-293T (435.3% increase, p= 0.0043) and HeLa (95.2% increase, p=0.002) cells compared to single transfection (Em_NPM1_mut only). This supports the tug-of-war hypothesis, proposing that the ratio of both wild-type and mutant constructs affect their localisation due to the formation of heterooligomers [28]. Curiously, we did not observe the characteristic nucleoli rim localisation of ectopic NPM1wt (Figures 2A, B) that was readily apparent with antibody staining (Figure 1B). We hypothesised that this discrepancy could either be a result of overexpression or due to fluorescent reporter (scarlet tag) interference. To test this, we used CRISPR-Cas9 methodology to edit the NPM1 gene, inserting a scarlet or emerald fluorescent protein, at the endogenous locus. Fluorescence imaging of the engineered cell lines, co-labelled with anti NPM1wt antibody, confirmed appropriate nucleoli rim localisation (Figure 2D yellows arrows). This serves as an important technical observation for the wider NPM1 field, demonstrating that NPM1 protein can tolerate reporter fusions, but only when expressed at endogenous, or near-endogenous levels.

**Figure 2.**
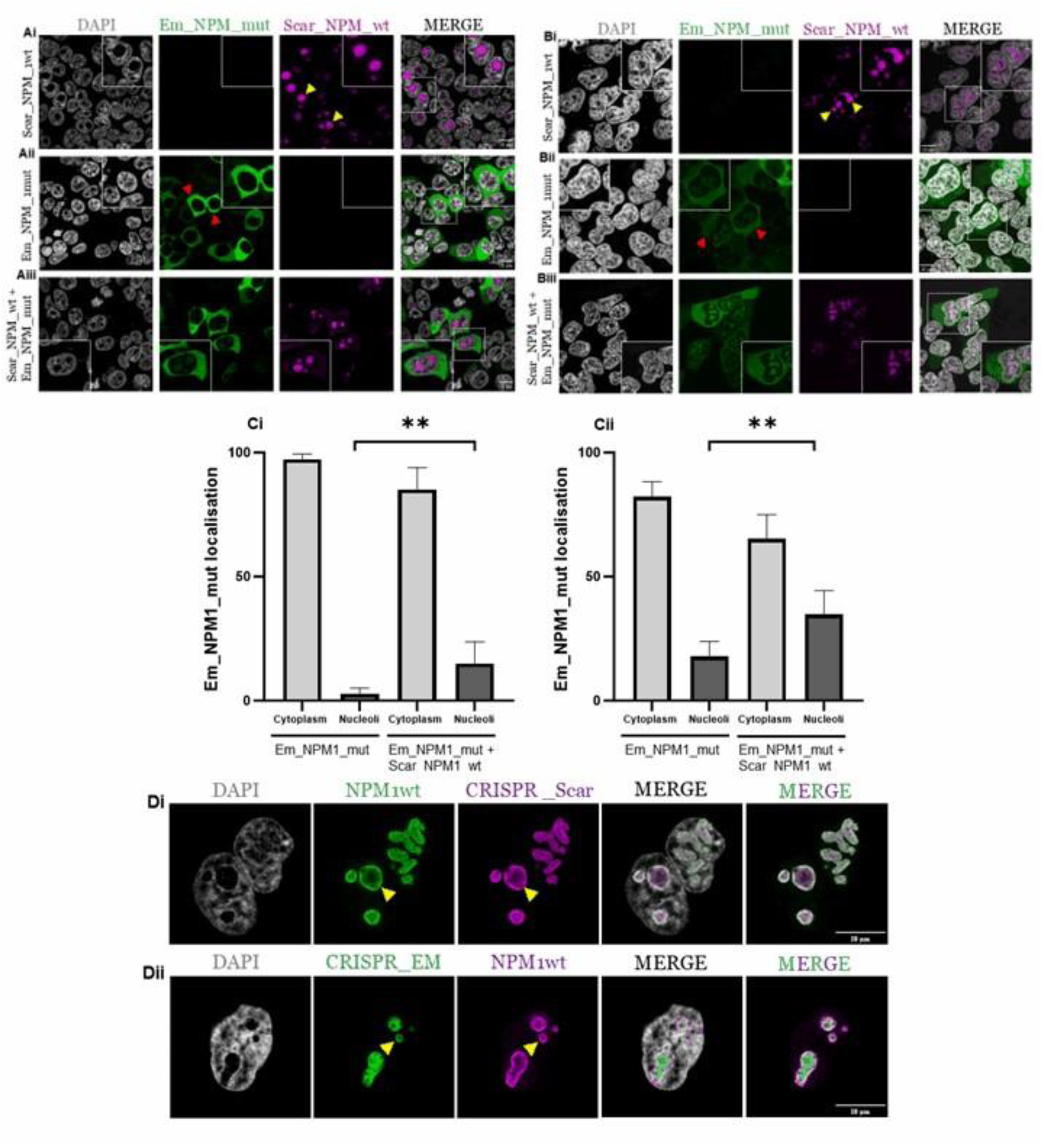
Subcellular localisation of wild-type and mutant NPM1 revealed by high-resolution confocal microscopy. **(A)** HEK-293T and **(B)** HeLa cells were transfected with Scar_NPM1_wt and Em_NPM1_mut constructs for 24 hrs. Yellow arrows indicate NPM1wt localisation in the nucleoli. Red arrows indicate NPM1 mutant localisation to the cytoplasm. **(C)** Subcellular localisation of Em_NPM1_mut in HEK-293T (i) and HeLa cells (ii) was quantified using Fiji. Columns, data from 20 representative cells: bars, SD **\*\***p=<0.005 (unpaired t-test). **(D)** Scarlet (Di) and Emerald (Dii) fluorescently tagged NPM1wt knock-in HeLa cells were created by CRISPR-Cas9 gene editing. Cells were assessed using either the fluorescent tag or stained using antibodies against NPM1wt. Yellow arrows indicate NPM1wt localisation to the nucleoli rim.

### NPM1-mutated cells lack a nucleoli rim and have distorted nucleoli compared to NPM1 wild-type cells

We next investigated the subcellular localisation and impact of NPM1wt and NPM1c+ on nucleolar architecture in leukaemia cell lines. We used OCI-AML3 and IMS-M2 cell lines, both of which are heterozygous for NPM1wt and NPM1c+. OCI-AML2 cells, expressing only NPM1wt, were used as a reference control.

A striking difference in nucleoli architecture was observed between the NPM1 mutated lines and the OCI-AML2 control cells (Figure 3A). In OCI-AML2 cells, imaging and line-scan analyses revealed that endogenous NPM1wt localised neatly to the nucleoli rim (Figure 3B – presence of double peaks) and the nucleoli exhibit typical healthy circular morphology (Figure 3C). In contrast, in OCI-AML3 and IMS-M2 cells, endogenous NPM1wt failed to localise to the nucleoli rim (Figure 3B – presence of single peaks) and instead showed a diffuse distribution within the remaining nucleolar compartment. In addition, the nucleoli in these cells exhibit irregular and abnormal nucleolar morphology (Figure 3C).

**Figure 3.**
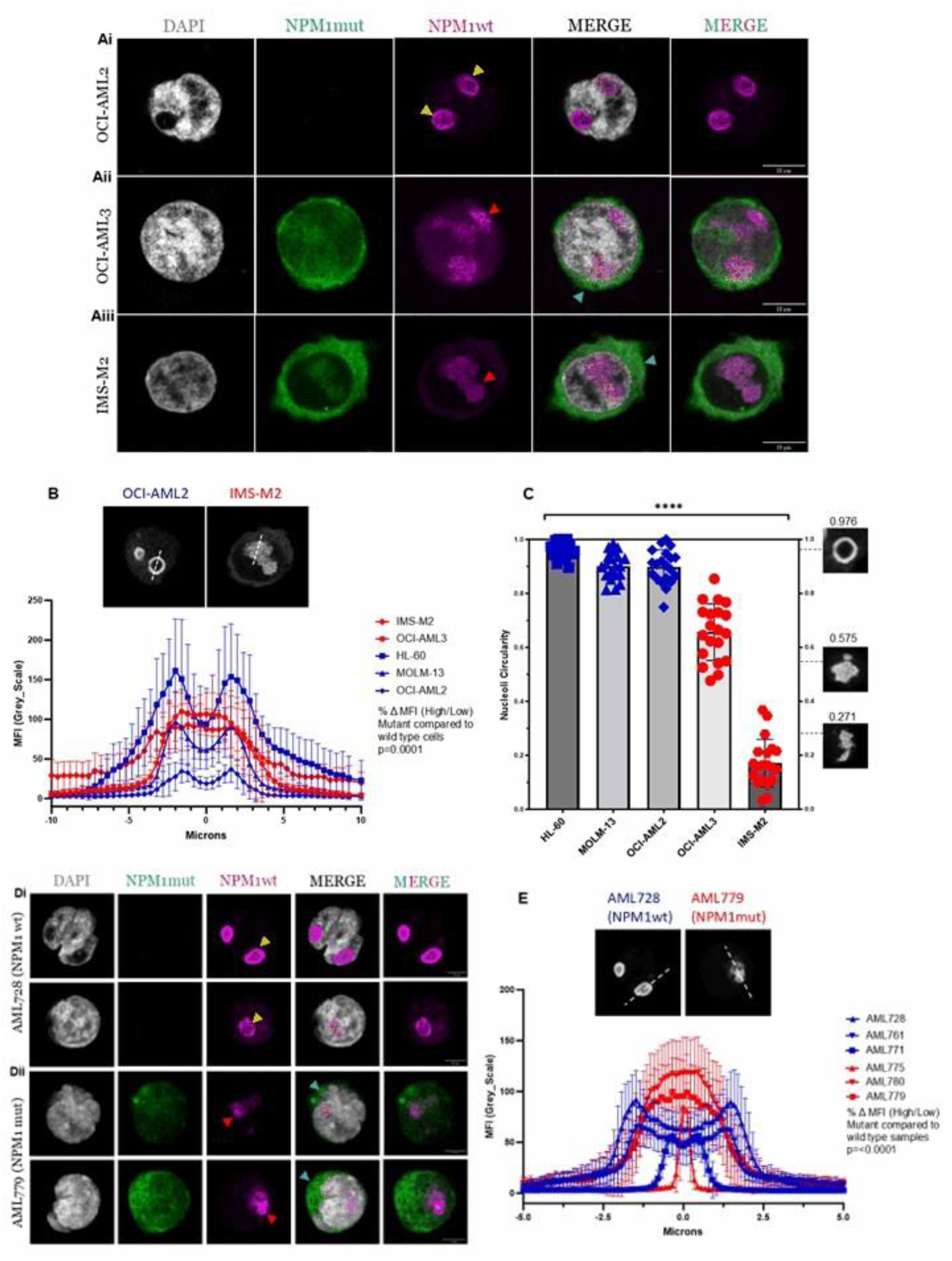
NPM1 mutated cells lack a nucleoli rim and have distorted nucleoli compared to NPM1 wild-type cells. **(A)** AML cell lines were stained with DAPI and antibodies against NPM1 mutant and NPM1wt. Yellow arrows indicate NPM1wt localisation to the nucleoli rim in NPM1 wild-type OCI-AML2 cells (Ai). Red arrows indicate a lack of nucleoli rim and distorted nucleoli in the OCI-AML3 (Aii) and IMS-M2 (Aiii) NPM1 mutated cells. Blue arrows indicate NPM1 mutant localisation to the cytoplasm. **(B)** NPM1wt localisation in the nucleolus of AML cell lines was quantified using the line scan tool in Fiji. NPM1 wild-type cell lines in blue, NPM1 mutant cell lines in red, mean of 20 nucleoli for each cell line: bars, SD, p=0.0001 (unpaired t-test). **(C)** AML cell line nucleoli circularity was measured using the threshold tool in Fiji. Columns, mean of 20nucleoli for each cell line: bars, SD ****p=<0.0001 (unpaired t-test). **(D)** Primary AML samples were stained with DAPI and antibodies against NPM1 mutant and NPM1wt. Yellow arrows indicate NPM1wt localisation to the nucleoli rim in NPM1 wild-type samples (Di). Red arrows indicate a lack of nucleoli rim and distorted nucleoli in the NPM1 mutated samples (Dii). Blue arrows indicate NPM1 mutant localisation to the cytoplasm. **(E)** NPM1wt localisation in the nucleolus of primary samples. NPM1 wild-type samples in blue, NPM1 mutant samples in red. Columns mean of 20 nucleoli for each sample: bars, SD p=0.0001 (unpaired t-test).

To further validate these findings, we imaged two additional NPM1wt AML cell lines, HL-60 and MOLM-13 (Supplementary Figure 1). Consistent with OCI-AML2 cells, endogenous NPM1 in these lines localised to the nucleoli rim and the nucleoli maintained a circular phenotype, unlike the diffuse NPM1 localisation and irregular nucleoli shape observed in the mutated cell lines. Quantitative analysis confirmed highly significant differences between NPM1wt and NPM1c+ cells in nucleolar rim intensity (p=0.0001) (Figure 3B) and nucleolar circularity (p<0.0001) (Figure 3C).

To extend these findings to a more clinically relevant context, we next examined six primary patient AML samples. In NPM1wt cells (n=3) NPM1 localised to the nucleoli rim and nucleoli appeared circular (Figure 3D). In contrast in NPM1c+ mutated samples (n=3), NPM1 was absent from the nucleoli rim, showing diffuse staining in the nucleolus, which appeared distorted (Figure 3D). Quantification of images, using pixel density line scans, confirmed a highly significant difference in nucleolar rim intensity between NPM1wt and NPM1c+ cells (p=<0.0001) (Figure 3E).

### Reversible induction of aberrant nucleolar architecture: The impact of NPM1c+ expression in wild-type AML cells and NPM1wt expression in mutated AML cells

To dissect the relationship between NPM1c+ expression and aberrant nucleolar architecture – particularly nucleolar rim integrity, we conducted reciprocal mutant/wt expression experiments exploiting the genetically distinct OCI-AML2 and OCI-AML3 cell lines. When NPM1c+ was expressed in the OCI-AML2 cells, it forced a shift from normal nucleolar architecture to an aberrant phenotype characterised by the loss of the nucleolar rim (Figure 4A, C). Conversely, the introduction of NPM1wt into the OCI-AML3 cells, restored nucleolar architecture and reformed the nucleolar rim (Figure 4B, C). The changes in NPM localisation, quantified via line scan analyses, were highly significant (p=0.006 and 0.005) (Figure 4C).

**Figure 4.**
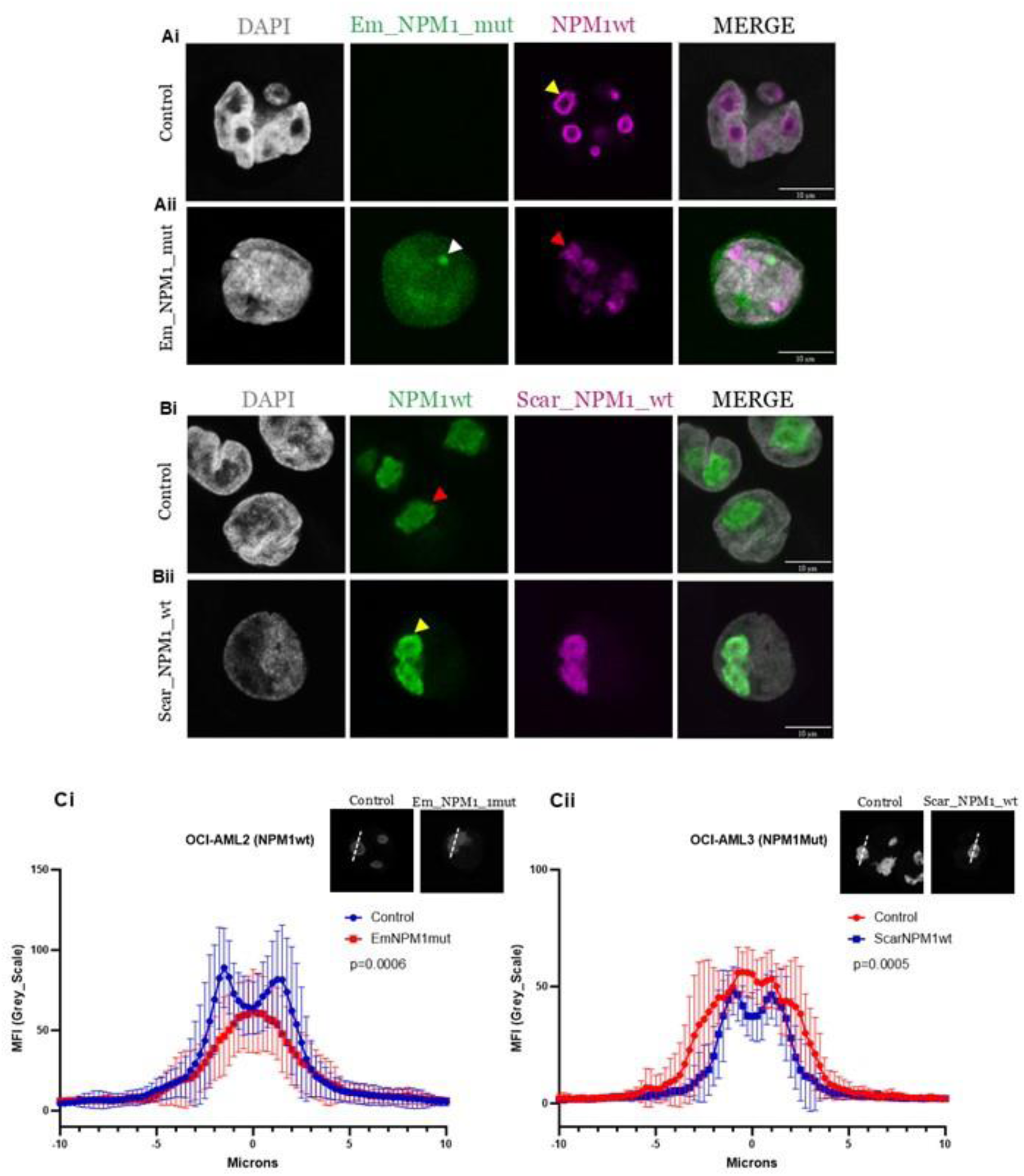
Reversible induction of aberrant nucleolar architecture: The impact of NPM1c+ expression in wild-type AML cells and NPM1wt expression in mutated AML cells. **(A)** OCI-AML2 and **(B)** OCI-AML3 cells were transfected with Em_NPM1_mut or Scar_NPM1_wt respectively for 24 hrs and stained with DAPI and NPM1wt antibody. Yellow arrows indicate NPM1wt nucleoli phenotype (Ai and Bii). Red arrows indicate NPM1 mutant nucleoli phenotype (Aii and Bi). White arrow indicates NPM1 mutant protein aggregate (Aii). **(C)** NPM1wt localisation at the nucleolus of OCI-AML2 cells (Ci) and OCI-AML3 cells (Cii), p=0.0006 and p=0.0005 (unpaired t-test). Mean of 20 nucleoli for each condition: bars, SD.

These findings indicate that loss of nucleolar integrity in NPM1 driven AML does not precede mutation of NPM1, but rather, that NPM1 is the initial cause – but importantly is also reversible.

Additionally, during our investigation, we discovered an unexpected novel feature of NPM1c+ mutants: following transfection of the Em_NPM1_mut construct into the OCI-AML2 cells, we consistently observed that a portion of NPM1c+ assembles into protein aggregates, (Figure 4Aii white arrow). These aggregates were not formed when Scar_NPM1_wt was introduced into OCI-AML3 cells.

### Mutant NPM1 protein forms distinct pools of protein aggregates

To investigate whether the protein aggregates formed by expressing Em_NPM1_mut were inherent to NPM1 mutated cells, or merely an artifact of overexpression, we re-examined our extensive collection of fluorescence images. Strikingly, we consistently observed NPM1 mutant protein aggregates, in the NPM1 mutated cell lines but also, importantly, in the patient samples (Supplementary Figure 2). The aggregates appear to represent a distinct pool of mutant NPM1 protein as they were not recognised by the NPM1wt specific antibody.

Furthermore, when we ectopically expressed the mutant NPM1 construct in HEK-293T and HeLa cells, we observed similar aggregate formation, which were absent when we ectopically expressed the NPM1wt construct in the same cell lines (Supplementary Figure 2). For the first time, we report that the majority of cells, naturally expressing NPM1c+ mutations contain NPM1c+ aggregates (OCI-AML3 85% and IMS-M2 90%), and that aggregates were seen in all cells of two of the primary samples analysed (Supplementary Figure 3). The number of aggregates per cell was 2.28 ±1.58 and 2.26 ±1.7 in the two NPM1 mutated cell lines and 4.1 ±1.8 and 2.45 ±1.19 in two of the NPM1 mutated primary samples. Curiously, we noticed that aggregates were not restricted solely to one compartment. 24-32.5% of aggregates were in the cytoplasm and 28-32.5% located in the nucleus, for both mutant cells lines (Figure 5A, 5B and 5C). Notably, a large fraction (28-43%) were located on the cytoplasmic interface of the nuclear membrane. Distribution of the aggregates in the NPM1 mutated primary samples differed slightly, with most of the aggregates (53-55%) localised to the nucleus (Supplementary Figure 3). Analogous to the cell lines, a large fraction appeared tethered to the cytoplasmic side of the nuclear membrane. The significance of this variability remains elusive but will be the subject of future study.

**Figure 5.**
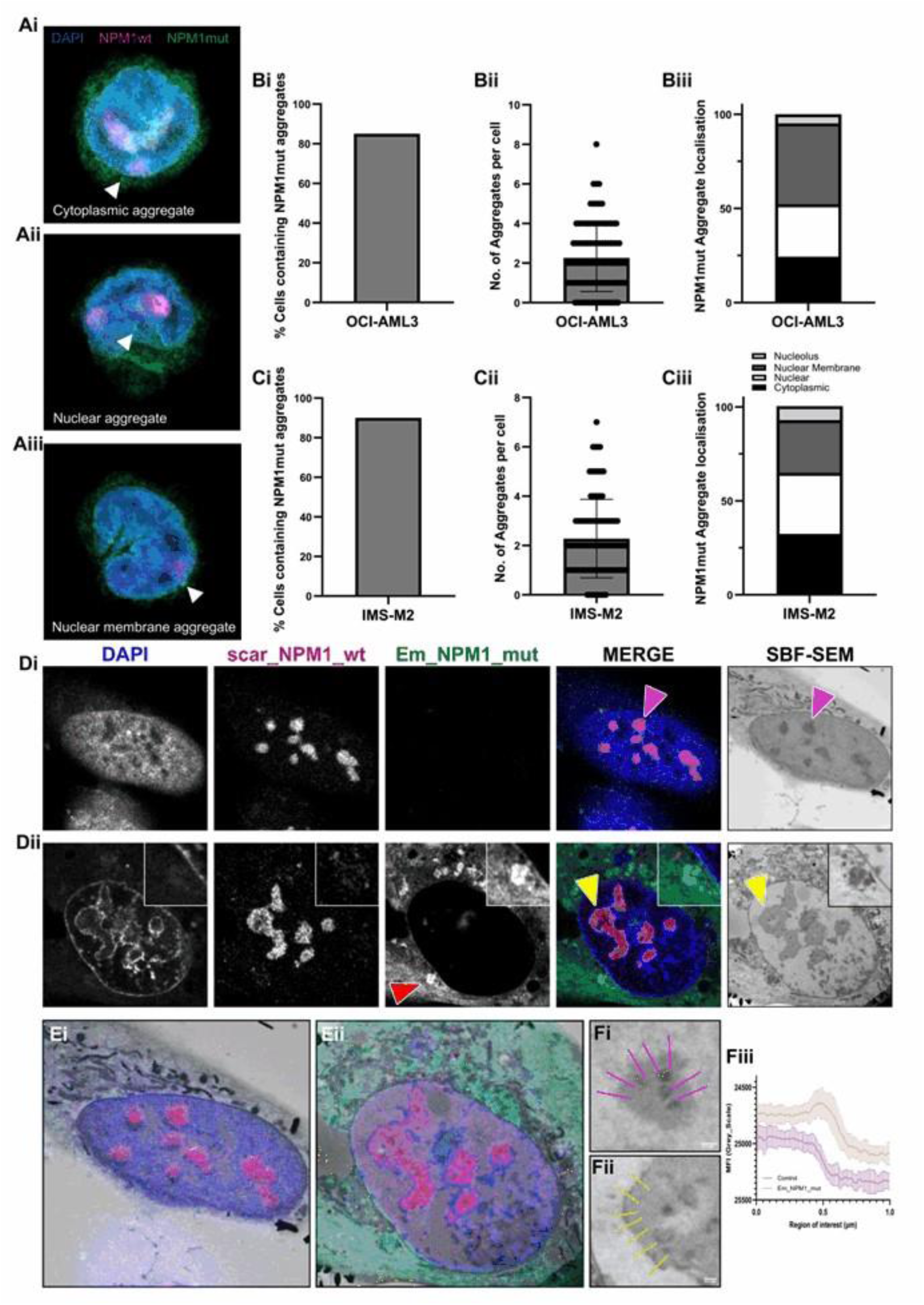
NPM1 mutant protein forms distinct pools of protein aggregates. **(A)** NPM1 mutated OCI-AML3 cells were stained with DAPI and antibodies against NPM1 mutant and NPM1wt. Images show representative examples of aggregate localisation (indicated by white arrows) in the cytoplasm, in the nucleus and at the nuclear membrane. Fluorescent image acquisition was performed using a ZEIS LSM 900 confocal microscope in Airyscan mode with a x100 oil immersion objective. **(B)** and **(C)** Aggregate quantification was performed by collating images from multiple slices along the Z-axis with a sampling depth of 2μm and analysed as maximum intensity projections. Columns, data from 100 representative cells: bars, SD. **(D)** Scarlet tagged NPM1wt knock-in HeLa cells were subjected to 24 hr Em_NPM1_mut transfection (Dii) or transfection control (Di). Light microscope images are shown in the first four panels with EM overlays in the far-right panels. Red arrow in Dii indicates an NPM1 mutant aggregate. Yellow arrows in Dii indicate an electron dense region surrounding the nucleoli that is not seen in control cells (pink arrows). **(E)** 3D modelling of EM images using Amira software demonstrates clear registration with the original LM images. **(F)** Electron density measured from the nucleolus to nucleoplasm in control cells (Fi) or Em_NPM1_mut expressing cells (Fii) was quantified using the line scan tool in Fiji (Fiii), bars, SD.

Given that most nucleolar components are dense enough to be visualised by electron microscopy (EM) – we next explored whether these mutant NPM1 aggregates could be observed at the ultra-structural level. We used SuperCLEM [29], an elegant multimodal imaging technique allowing rare events (such as aggregates) to be captured using both super resolution light microscopy (LM) and EM.

We identified and examined cells of interest - those expressing only NPM1wt as controls (Figure 5Di) and those expressing Em_NPM1_mut (Figure 5Dii). Aggregates were clearly visible in the cells expressing mutant NPM1 (red arrow). We prepared these for SuperCLEM and revisited our chosen cells using serial block face scanning electron microscopy (SBF-SEM) (Figure 5D right panels). Clear registration could be achieved between LM and EM images, confirming re-location of not only the same cells, but same area of each cell (Figure 5E). SuperCLEM revealed several structural differences between control and mutant NPM1 expressing cells. 1) Registration of optical (light) and physical (EM) sections allowed us to visualise areas of the cell corresponding to aggregates (Figure 5Dii red arrow). Clear, electron dense structures were present at sites of registered aggregate sites (Figure 5Dii inset zoom). 2) There was a clear shift in the electron density of the nucleoplasm, between control and NPM1 mutant samples. 3) Linked to point 2, in the Em_NPM1_mut expressing cells we observed the presence of a density surrounding the nucleolus (Figure 5Dii far right panels - yellow arrows) which was not seen in the control cells (Figure 5Di far right panels - pink arrows). This was quantifiable using a line scan analysis measuring the pixel densities between the nucleolar body, through the rim and into the nucleoplasm (Figure 5F). The presence of the “shoulder” in Figure 5Fiii reflects the electron dense region.

## Discussion

Mutations in exon 12 of the *NPM1* gene were first identified almost 20 years ago and represent one of the most common genetic alterations in AML [3]. The mutations are always heterozygous, suggestive of an essential role for wild type NPM1, although preferential transcription of the mutant allele has been reported [30]. The NPM1c+ mutation has important clinical and prognostic implications, and the focus of much research has centred on the mis-localisation of the mutated protein from the nucleolus to the cytoplasm [31, 32]. Here we report that NPM1wt subcellular localisation is also greatly impacted in NPM1 mutated cells. Specifically, NPM1 dissociates from its usual location at the nucleoli rim, and instead exhibits a diffuse staining pattern in the nucleoli with resultant breakdown of nucleoli structure. We also report that the aberrant nucleoli phenotype in NPM1 mutated cells is reversible and conversely that aberrant nucleoli phenotype can be induced in NPM1wt cells. To our knowledge this is the first demonstration of this kind of nucleoli architecture manipulation in AML cells but is of significance as we search for new strategies to treat NPM1 driven AML, including the potential of nucleoli as vulnerable targets. Morphologically, nucleoli consist of three distinguishable phase separated regions, the fibrillar centre, the dense fibrillar component and the granular component. More recently, a fourth nucleoli sub-compartment, the nucleoli rim, has been identified. We demonstrate that under normal physiological conditions NPM1 locates to the nucleoli rim, but in NPM1 mutated cells this rim localisation is lost, resulting in distorted nucleoli. One caveat to this is that we did not see NPM1 localisation to the nucleoli rim when we ectopically expressed NPM1 in HeLa and HEK-293T cells. We therefore suggest that care should be taken when interpreting data from cells where NPM1 has been overexpressed. The localisation of NPM1 to the nucleoli rim is seemingly dependent on the amount of NPM1wt in the nucleolus, as we report that both knockdown of NPM1wt in NPM1wt cells, and mis-localisation of a portion of NPM1wt to the cytoplasm in NPM1c+ cells, result in nucleoli distortion. Studies have implicated NPM1 in maintaining a normal nucleolar structure through its interaction with ribosomal subunit precursors, and cells depleted of NPM1, exhibit deformed nucleoli and rearrangement of peri-nucleolar heterochromatin. Indeed, our ultrastructural analysis, using SuperCLEM, may support this, as the expression of NPM1C+ mutant caused apparent changes in chromatin organisation observable by both super resolution light microscopy (DAPI staining) but also at the underlying EM level, via an electron dense region surrounding the nucleolus, typically reminiscent of heterochromatin staining. Disruption of the nucleolus and subsequent dysregulated ribosome biogenesis has been linked to tumour initiation and cancer progression implicating the nucleoli as a potential therapeutic approach [33]. In fact, it has been suggested that the nucleoli of NPM1 mutated AML cells might be particularly susceptible to drugs that induce a nucleolar stress response due to the partial depletion of NPM1 caused by both haploinsufficiency and retention of a fraction of NPM1wt by NPM1c+ in the cytoplasm [34]. We have previously reported that etoposide or cytarabine induced DNA damage alters the subcellular localisation of mutated NPM1 back to a predominantly nucleolar distribution [35]. Thus, the induction of nucleolar stress has emerged as a potential therapeutic strategy for NPM1 mutated AML.

The N-terminus of NPM1 contains a self-oligomerization core that is responsible for the formation of NPM1 pentamers and the oligomerization state of NPM1 is critical for its normal biological functions [36, 37]. It is therefore important to explore the molecular links between oligomerization and NPM1 driven leukaemia. Our high-resolution imaging consistently revealed the presence of distinct pools of NPM1 mutant protein aggregates in both NPM1 mutated cell lines and primary samples. Aggregates were also present when we ectopically expressed mutant NPM1 in HEK-293T and HeLa cells, but not with NPM1wt ectopic expression, suggesting that the mutated C-terminal of NPM1c+ is responsible for aggregate formation. The significance of the variability in the localisation of the aggregates remains unclear but is likely related to how and when they are assembled and warrants further study.

Our Super-CLEM analysis revealed that the aggregates co-register to electron dense areas. These could correspond to self-assembling phase separated compartments or could be a result of abnormal nucleolar disassembly. EM also revealed, a redistribution of chromatin and recruitment of an electron dense nucleoli surface, potentially indicative of a change in chromatin from a more open (euchromatic) to a more closed (heterochromatic) state, consistent with reported disruption to gene expression profiles. Interestingly, by means of complementary biophysical techniques, studies have linked an aggregation propensity of distinct regions of the improperly folded C-terminal domain to leukaemogenesis in NPM1 mutated AML. By comparing the conformational and aggregate forming ability of the entire C-terminal domains of NPM1wt and NPM1c+ the authors reported that only the C-terminus on NPM1c+ and not NPM1wt can form amyloid-like aggregate assemblies [38, 39].

Here, for the first time, we report NPM1 mutant protein aggregate formation and a distinct nucleoli phenotype in NPM1 mutated AML cells both of which represent potential therapeutic vulnerabilities. The fact that this particular pool of mutant NPM1 cannot readily form multimers with NPM1wt is of critical importance, therefore the spatiotemporal origins (and disassembly) and biochemical profiles of these aggregates need to be carefully studied in future studies.

## Materials and Methods

### Cells

OCI-AML3, OCI-AML2, HL-60 and MOLM-13 cells were obtained from the Leibniz Institute (DSMZ, Braunschweig, Germany). IMS-M2 cells were a kind gift from Carolien Woolthuis, University of Groningen, Netherlands. Cells were maintained in RPMI 1640 medium with 10% foetal calf serum (Fisher Scientific, Hampton, New Hampshire, USA) and 2mM L-glutamine (Sigma-Aldrich, St. Louis, Missouri, USA). HeLa and HEK-293T cells were obtained from DSMZ and maintained in DMEM with 10% FCS. Cultures were sustained at 37°C in 5% CO_2_ and all experiments were performed with cell lines in log phase. Monthly mycoplasma testing was performed using PlasmoTest mycoplasma contamination detection kit (InvivoGen, Toulouse, France). AML patient samples were obtained with informed consent in accordance with national and international guidelines and approval by the authors’ institutional review board (REC# 06/Q2403/16). Mononuclear cells were obtained by standard density gradient centrifugation from bone marrow or peripheral blood samples and cells were cryopreserved until use. The presence of NPM mutations were identified as previously described [40].

### Constructs

Gene inserts were synthesised by GeneArt (Regensberg, Germany) as follows: For the NPM1wt insert, unique SacI (5’) and EcoRI (3’) sites were added to full length hsNPM1wt (NM_002520.7). For the NPM1 mutant insert, the C-terminal of the NPM1wt insert was replaced with the mutated sequence (gat ctc tgt ctg gca gtg gag gaa gtc tct tta aga aaa tag). NPM1wt and NPM1 mutant inserts were cloned into the SacI and EcoRI sites of pmScarlet_C1 (Addgene, Watertown, Massachusetts, USA #85042) and mEmerald-C1 (Addgene, #53975) plasmids respectively by standard methods.

### Transfections

HeLa or HEK-293T cells in exponential growth were seeded in twelve-well plates on glass coverslips and cultured overnight. Transfections were performed using Polyplus jetPRIME (VWR, Lutterworth, Leicestershire, UK) with either 0.8µg of Scar_NPM_wt or Em_NPM_mut constructs. NPM1 wild-type OCI-AML2 and NPM1 mutant OCI-AML3 cells were nucleofected with 1µg Em_NPM_mut or Scar_NPM_wt constructs respectively using Nucleofector Kit-T (cat# VCA-1002, Lonza, Basel, Switzerland) with programme X-001 using a Nucleofector-2b device (Lonza). Construct expression and ectopic protein localisation was confirmed using fluorescence microscopy.

### Generation of fluorescently tagged NPM1 HeLa cells using CRISPR/Cas9-mediated gene editing

DNA oligonucleotides (Forward 5’ ACATGGACATGAGCCCCCTG 3’, Reverse 5’ GATAGTTCTGGGGCCTCAGG 3’) used for gRNA synthesis were designed using the Benchling CRISPR gRNA Design Tool available at (www.benchling.com, accessed on 15 May 2022) and ordered through Sigma. gRNA was cloned into the Bbs1 restriction site of the pX330-U6-Chimeric_BB-CBh-hSpCAas9-hGem (1/110) vector (Addgene #71707) before transformation into NEB^®^ 5-alpha Competent *E. coli* (New England Biolabs, Ipswich, Massachusetts, USA). DNA was prepared from single colonies before sequence verification using an Applied Biosystems^™^ (Waltham, Massachusetts, USA) 3130*xl* genetic analyser. Homology repair plasmids (HDR) designed to introduce mScarlet or mEmerald to the N-terminus of NPM1 using a pUC18 backbone were ordered from GenScript (Piscataway, New Jersey, USA). HeLa cells were nucleofected with the HDR plasmid and the gRNA cas9 plasmid and allowed to recover for 3 days before selection with G418 for 7 days. To remove antibiotic selection via the loxP sites, and allow transcription of tagged protein, cells were transfected with pMSCVpuro-Cre plasmid (Addgene #34564). Cells were then treated with puromycin (Fisher Scientific) for 3 days and single cell sorted into ninety-six well plates. Clones were screened by PCR genotyping.

### RNA Interference

For RNAi treatments HeLa or HEK-293T cells in exponential growth were seeded in twelve-well plates on glass coverslips and cultured overnight. Transfections were performed using Polyplus jetPRIME with 15nM siRNAs targeting both the wild-type and mutant NPM1 (AAAGGTGGTTCTCTTCCCAAA) (Hs_NPM1_7, SI02654960, Qiagen, Hilden, Germany) or AllStars negative control siRNA (cat# 1027281, Qiagen). Following siRNA interference NPM1 knockdown was confirmed by western blot and indirect immunofluorescence. For the rescue experiments HEK-293T cells at 50% confluence were transfected with 15nM NPM1 targeting siRNA (Hs_NPM1_7) or AllStars negative control siRNA. Following 2 days siRNA interference cells were transfected with 0.8µg of Scar_NPM_wt construct and expressed for 24 hrs before validation by indirect immunofluorescence.

### Indirect immunofluorescence and microscopy

Primary antibodies were used as follows: NPM1wt (Mouse monoclonal, #32-5200 (Thermo fisher, Waltham, Massachusetts, USA) 1:60; NPM1 mutant (Rabbit polyclonal, Thermo fisher #PA1-46356) 1:60; Nucleolin (Rabbit polyclonal, Abcam, Cambridge, UK, #ab22758) 1:100. Fluorescence-labelled secondary antibodies were applied at 1:400 (Jackson ImmunoResearch, Ely, UK). For immunofluorescence, cells were fixed in 3.5% paraformaldehyde for 15 min, permeabilized in 0.3% Triton X-100 for 5 min and blocked in 2% blocking solution (Thermo fisher). Cells were incubated overnight with the primary antibodies washed in PBS and secondary antibodies were applied for 1 hr before counter-staining with DAPI. Fluorescent image acquisition was performed using a Leica (Wetzlar, Germany) TCS SPE confocal microscope with a x63 oil immersion objective. Images were exported as TIFF files and imported into Fiji for final presentation [41].

### Image analysis

NPM1 localisation in the nucleolus was quantified using the line scan tool in Fiji. Nucleoli circularity was determined using the threshold tool. Aggregate quantification was performed by collating images from multiple slices along the Z-axis with a sampling depth of 2μm and presented as maximum intensity projections. For EM line scan: 5-10 1μm line scans were used per nucleoli (4-8 nucleoli in total). Pixel densities were retrieved with the line scan origin located in the nucleolus and terminating in the nucleoplasm, with the nucleolar-to-nucleoplasm interface being approximately mid-line scan (0.5um). Pixel densities were recorded every 10nm.

### Western blot analysis

HeLa cells were subjected to 3 or 6 days NPM1 siRNA interference. Cell lysates were prepared, separated by sodium dodecyl sulphate polyacrylamide gel electrophoresis, and transferred to nitrocellulose membranes. Primary detection antibodies were Mouse anti-NPM1 (1:1000) (Thermo fisher #32-5200), Rabbit anti-Nucleolin (1:1000) (ab22758), Rabbit monoclonal anti-Actin (1:5000) (ab179467), and Mouse monoclonal anti-Actin (1:1000) (ab8226) from Abcam. Secondary detection antibodies were Donkey anti-Mouse 680 (#926-68072) and Donkey anti-Rabbit 680 (#926-68073) from LI-COR (Lincoln, Nebraska, USA). Band quantification was determined using Fiji.

### Super 3D Correlative Light and Electron Microscopy (Super-CLEM)

Scarlet tagged NPM1wt knock-in HeLa cells were seeded onto glass-bottomed, gridded dishes (MatTek Corporation, Ashland, Massachusetts, USA) and transfected with Em_NPM_mut construct or transfection control. Following a 24 hrs expression period, cells of interest were identified using a Zeiss (Oberkochen, Germany) LSM900 confocal microscope mounted with Airyscan module. Cells were located and their position mapped using transmitted light to visualise reference coordinates. The Super-CLEM processing method was an adapted version of a previously established protocol [42]. Images were acquired using a Gatan 3View serial block face system (Gatan, Pleasanton, California, USA) installed on a FEI Quanta 250 FEG scanning electron microscope (FEI Company, Hillsboro, Oregon, USA). Images were collected at a magnification of 5.7k, voxel size of 12 x 12 x 60nm and chamber pressure of 70 Pa at 4 kV. Post acquisition analysis was performed using Amira software (Thermo Fisher).

### Calculations and Statistics

Unpaired t tests were performed using GraphPad Prism version 10.0.2 for Windows, GraphPad Software, Boston, Massachusetts, USA, www.graphpad.com. P values of ≤0.05 were considered to represent significance.

---

**Supplementary Figure 1.**
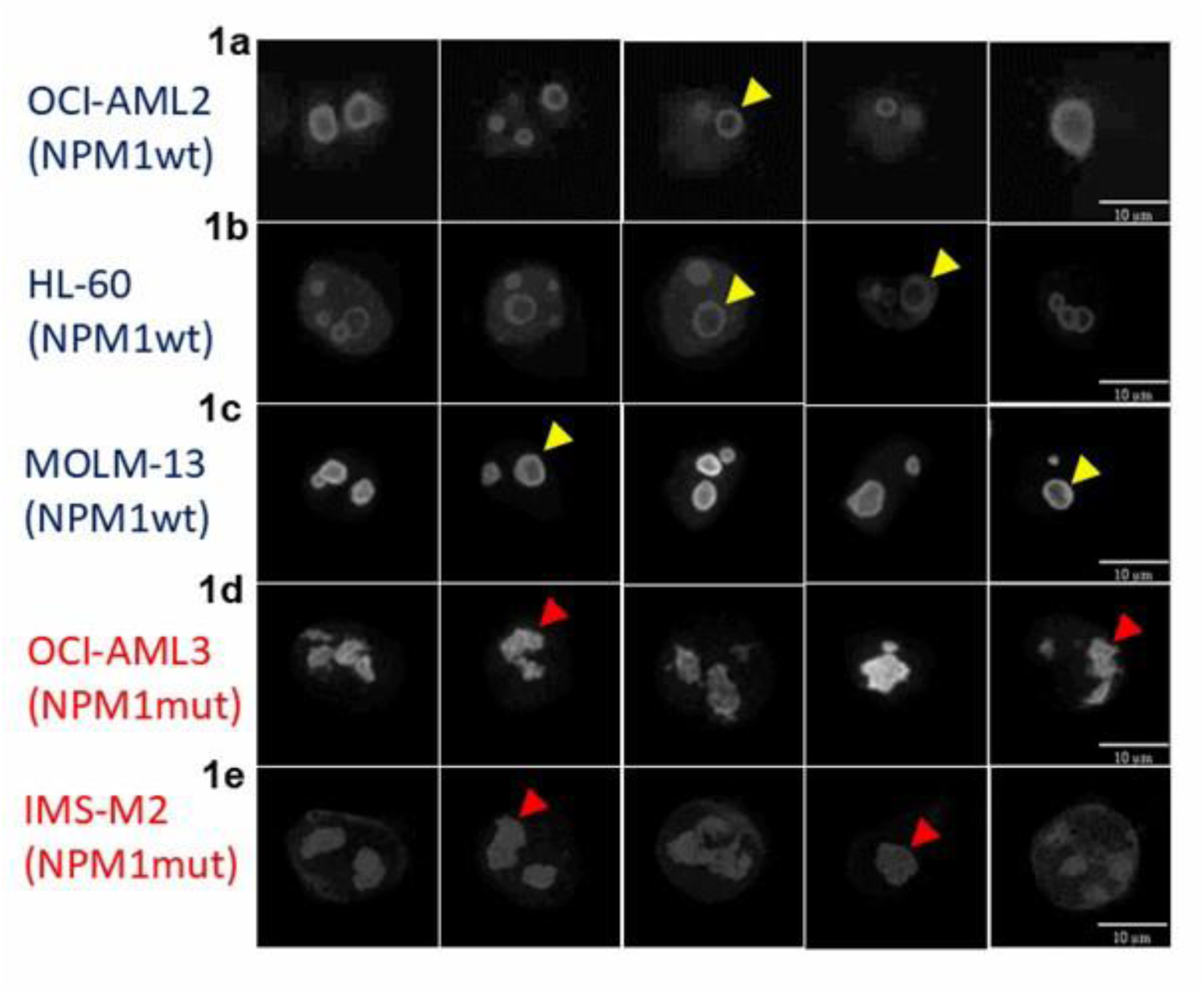
NPM1 mutated cells lack a nucleoli rim and have distorted nucleoli compared to NPM1 wild-type cells. AML cell lines were stained using an antibody against NPM1wt. Yellow arrows indicate NPM1wt localisation to the nucleoli rim in 3 NPM1 wild-type cell lines (1a, 1b and 1c). Red arrows indicate a lack of nucleoli rim and distorted nucleoli in NPM1 mutated OCI-AML3 and IMS-M2 cell lines (1d and 1e).

**Supplementary Figure 2.**
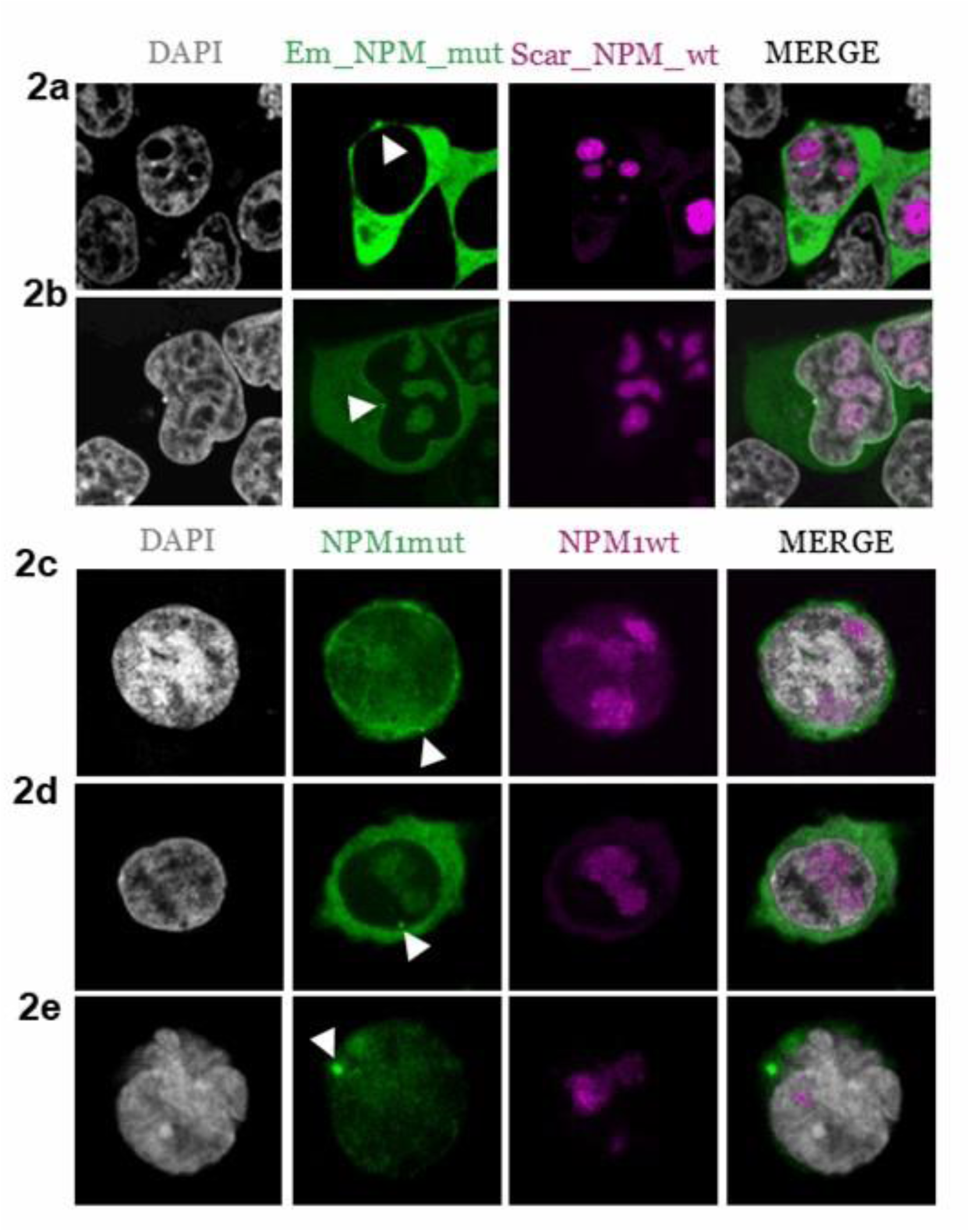
NPM1 mutant protein forms distinct pools of protein aggregates. **(2a)** HEK-293T and **(2b)** HeLa cells were transfected with Scar_NPM1_wt and Em_NPM1_mut constructs for 24 hrs. **(2c)** OCI-AML3 **(2d)** IMS-M2 or **(2e)** primary cells were stained with DAPI and antibodies against NPM1 mutant and NPM1wt. White arrow indicates NPM1 mutant protein aggregates.

**Supplementary Figure 3.**
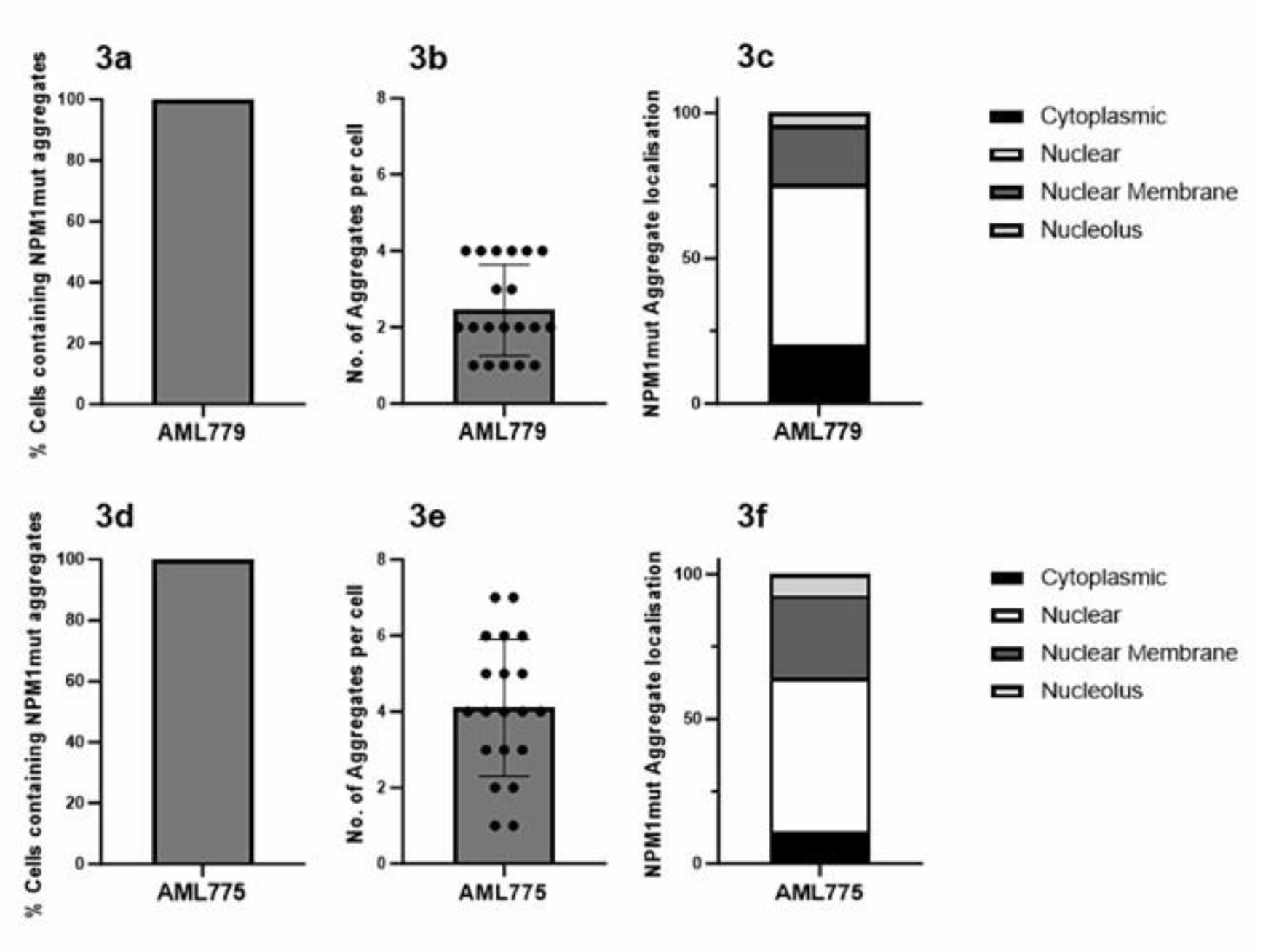
NPM1 mutant protein forms distinct pools of protein aggregates – Primary sample quantification. NPM1 mutated primary samples were stained with DAPI and antibodies against NPM1 mutant and NPM1wt. Aggregate quantification was performed by collating images from multiple slices along the Z-axis with a sampling depth of 2μm and analysed as maximum intensity projections. Columns, data from 20 representative cells: bars, SD.

